# Retron Editing for Precise Genome Editing without Exogenous Donor DNA in Human Cells

**DOI:** 10.1101/2021.05.11.443596

**Authors:** Xiangfeng Kong, Zikang Wang, Yingsi Zhou, Xing Wang, Linyu Shi, Hui Yang

**Affiliations:** HUIGENE Therapeutics Inc., Shanghai 200131, China; Institute of Neuroscience, State Key Laboratory of Neuroscience, Key Laboratory of Primate Neurobiology, Center for Excellence in Brain Science and Intelligence Technology, Chinese Academy of Sciences, Shanghai 200031, China; Shanghai Center for Brain Science and Brain-Inspired Intelligence Technology, Shanghai 201210, China

**Author notes:** These authors contributed equally to this work (Xiangfeng Kong, Zikang Wang). Correspondence to: Hui Yang.

**Keywords:** CRISPR, Retron, retron-ncRNA gRNA (rgRNA), genome editing

## Abstract

CRISPR-Cas9 mediated seamless genome editing can be achieved by incorporating donor DNA into the CRISPR-Cas9 target loci via homology-directed repair (HDR), albeit with relative low efficiency due to the inefficient delivery of exogenous DNA. Retrons are bacterial genetic element composed of a non-coding RNA (ncRNA) and reverse transcriptase (RT). Retrons coupled with CRISPR-Cas9 have been shown to enhance precise genome editing via HDR in yeast through fusing guide RNA (gRNA) to the 3’ end of retron ncRNA, producing multicopy single-stranded DNA (msDNA) covalently tethered to gRNA. Here, we further engineered retrons by fusing Cas9 with *E.coli* RT from different clades and joining gRNA at the 5’ end of retron ncRNA, and found that retron editing can achieve precise genome editing efficiently in human cells. By co-expression of Cas9-RT fusions and retron-ncRNA gRNA (rgRNA) in HEK293T cells, we demonstrated the rates of retron editing at endogenous genomic loci was up to 10 %. We expect our retron editing system could aid in advancing the *ex vivo* and *in vivo* therapeutic applications of retron.

## Introduction

Introducing precise modification into genome is of great significance in life science research, therapeutic application and agricultural development. The advent of facile RNA-programmable CRISPR-Cas system had dramatically advanced the development of precise genome editing (Anzalone et al., 2020; Cox et al., 2015; Doudna, 2020; Gao, 2021; Mao et al., 2019). CRISPR-Cas9 modifies the genome by introducing insertions and deletions (indels) via nonhomologous end-joining pathway (NHEJ), or homology-directed repair (HDR) at the target site after Cas9 inducing DNA double-strand breaks (DSB) (Cong et al., 2013; Yang et al., 2013). Theoretically, HDR can write the most extensive kind of modifications into the genome. Whereas, CRISPR-Cas9 stimulated DSB is predominantly repaired by NHEJ rather than HDR. In recent years, efforts have been made to enhance the HDR induced by Cas9-mediated DSB (Chu et al., 2015; Lin et al., 2014; Maruyama et al., 2015; Paquet et al., 2016; Richardson et al., 2016). Evidences have shown enhanced HDR by increasing the local abundance of donor DNA (Aird et al., 2018; Ma et al., 2017; Ma et al., 2020; Savic et al., 2018), indicating that the availability of donor DNA may be the rate-limiting step. In addition, the repulsion of cell membrane against exogenous DNA makes HDR-mediated precise genome editing a tough task due to the inefficient exogenous delivery of homologous donor DNA served as a template.

Retrons are bacterial phage-defense related operons composed of a specialized reverse transcriptase (RT) and a relevant non-coding RNA (ncRNA) which can be partially reverse transcribed by RT initiating at a conserved guanosine (G) residue to produce a multicopy single-stranded DNA (msDNA) (Gao et al., 2020; Millman et al., 2020; Yee et al., 1984). After being reverse transcribed, the msDNA is usually covalently tethered to the ncRNA through the 2’, 5’-phosphodiester bond between the priming G in ncRNA and 5’ end of msDNA (Dhundale et al., 1987; Lampson et al., 1989) (Fig. 1A). The reverse transcription process, of which the specialized RT recognizes the unique secondary structure of retron ncRNA, is highly specific (Hsu et al., 1989). Additionally, desired msDNA can be generated *in vivo* by replacing the dispensable region of retron ncRNA with desired sequences (Farzadfard and Lu, 2014; Inouye et al., 1997; Lopez et al., 2021; Mao et al., 1995). Therefore, retrons are promising biological sources for *in vivo* generation of DNA donors for HDR-mediated precise genome editing.

**Figure 1.**
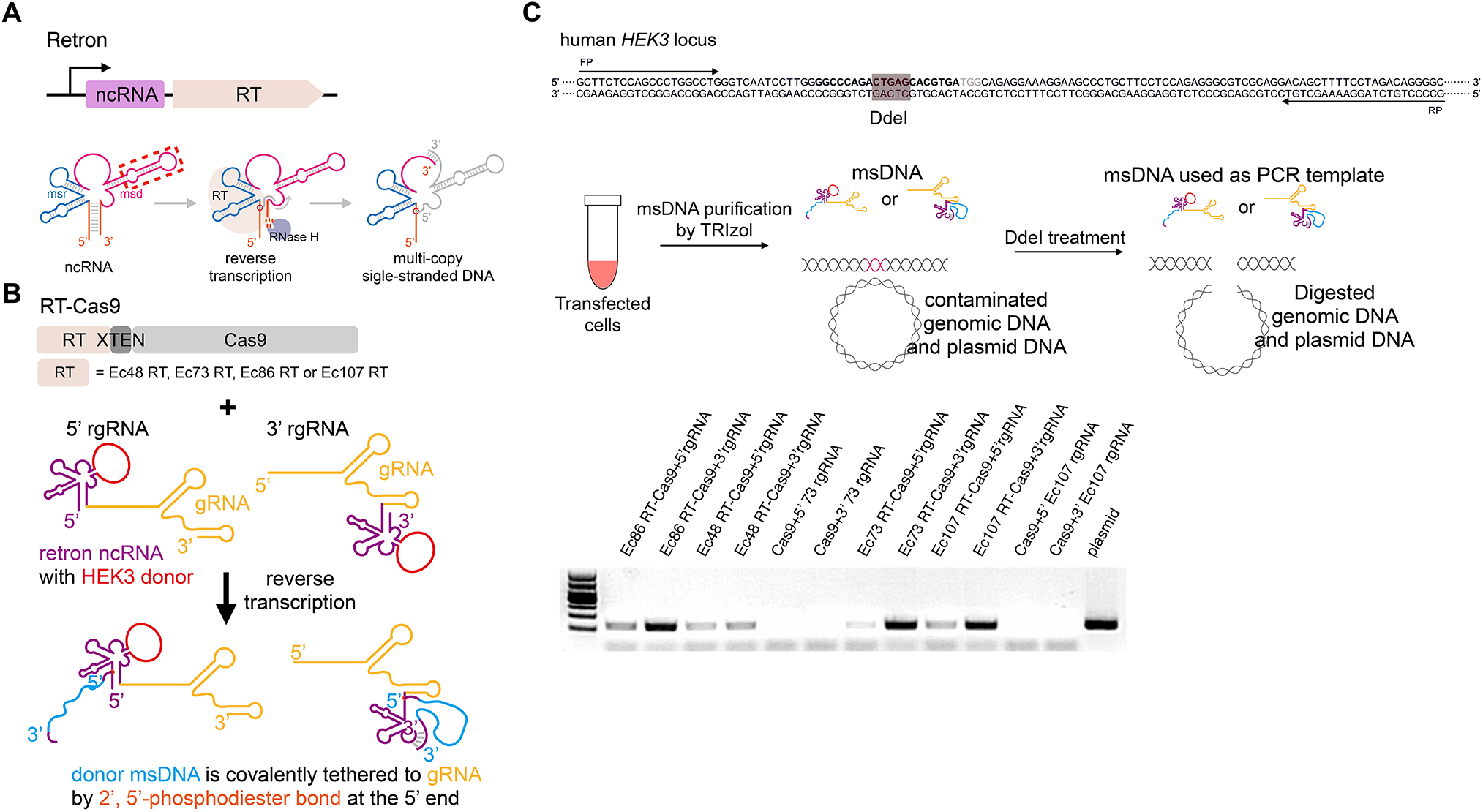
Retron-mediated generation of msDNA in human cells. **(A)** Schematic representing retron-mediated msDNA production. Retrons are composed of non-coding RNA (ncRNA) and a specific reverse transcriptase (RT). The msr and msd region of retron ncRNA form secondary structure which can be specifically recognized by relevant retron RT reverse transcribing part of msd region, generating multicopy single-stranded DNA (msDNA). After reverse trancription, the msd template is degraded by RNase H. The replaceable region of ncRNA, in which the donor sequence is inserted, is marked by red-dotted box. Red circle indicates 2’, 5’-phosphodiester bond between 2’ end of priming guanosine (G) and 5’ end of the msDNA. **(B)** Plasmids construction strategies for retron-mediated generation of msDNA in human cells. Human codon-optimized RT-XTEN-spCas9 fusions are driven by the CMV promoter, whereas the 5’ extended retron-ncRNA gRNA (5’ rgRNA), in which the retron ncRNA is fused to the 5’ of gRNA, or 3’rgRNA, is driven by the EF1⍺ promoter. Four E.coli retrons (Ec48 RT, Ec73 RT, Ec86 RT and Ec107 RT) from different clades were used in our retron editing system. The replaceable region of retron ncRNA was replaced by a 122nt modified HEK3 sequencce. **(C)** Determination of msDNA level. The msDNA abundance was determined by PCR. PCR was conducted using the DdeI digested product as template.

Recently, several studies have shown that when coupled with CRISPR-Cas9, retrons can be harnessed for highly efficient genome editing by *in vivo* producing donor single stranded DNA (ssDNA) in bacterium and yeast (Lim et al., 2020; Schubert et al., 2021; Sharon et al., 2018). Particularly, the retron-mediated reverse transcription has been described in mammalian cell (mouse NIH3T3 cell) (Mirochnitchenko et al., 1994). Therefore, it is plausible to assume the CRISPR-Retron system is suitable for precise genome editing in human cells. In this study, we detected the retron-mediated expression of ssDNA of interest in human cells. Moreover, we also show CRISPR-Retron mediated precise editing in human cells.

## Results

### Retron-mediated single-stranded DNA expression in human cells

Given that HDR-mediated precise editing can be enhanced by increasing the local abundance of donor DNA, and CRISPEY strategy showed highly efficient editing in yeast genome by *in situ* expressing retron-gRNA chimeric molecule (Sharon et al., 2018). Therefore, we attempted to *in situ* reverse transcribe the donor msDNA covalently tethered to gRNA by fusing retron ncRNA at the 5’ or 3’ end of gRNA, termed as the retron-ncRNA gRNA (rgRNA) (Fig.1B). Four experimentally validated *E.coli* retrons were evaluated for *in vivo* production of msDNA in human cells (Fig. 1B and S1). Additionally, we fused retron RT to the amino terminus or carboxy terminus of Cas9 with XTEN linker to increase the spatial proximity between RT and Cas9, which may enhance the retron editing in human cells by increasing the abundance of donor msDNA in the vicinity of DSB stimulated by Cas9 (Fig. 1B).

We first sought to test the relative abundance of msDNA in human cells. We found all of four selected RT enabled the expression of msDNA in human cells, and retron RT combining with 3’ rgRNA showed higher expression of msDNA comparing to that with 5’rgRNA (Fig. 1C). The results indicated that retron RT were reverse transcription-functional in human cells, inspiring us to further study the potential of retron editing at human endogenous genomic loci.

### Retron editing at endogenous genomic loci in human cells

Next, we investigated the potential of retron editing in human HEK293T cells (Fig. 2 and S2). We transfected the HEK293T cells with plasmids expressing different Cas9-RT fusions targeting *EMX1* and *HEK3*, and plasmids expressing 5’ rgRNA or 3’ rgRNA that can produce msDNA with homology donor sequences as well (Fig. 2A and 2B). To determine the retron editing efficiency at *EMX1* and *HEK3*, we cloned and Sanger-sequenced genomic DNA PCR amplicons of the target region using blunt-end cloning. Among four of retron tested, Ec86, Ec73 and Ec107 achieved varying degrees of precise editing and Ec73 combining with CRISPR-Cas9 showed highest activity, up to 10% (Fig. 2C). Restriction-fragment length polymorphism (RFLP) results confirmed Sanger-sequencing results (Fig. S2A), indicating that Ec73 RT-Cas9 fusion combining with 3’ Ec73 rgRNA can be harnessed for efficient precise genome editing in human cells.

**Figure 2.**
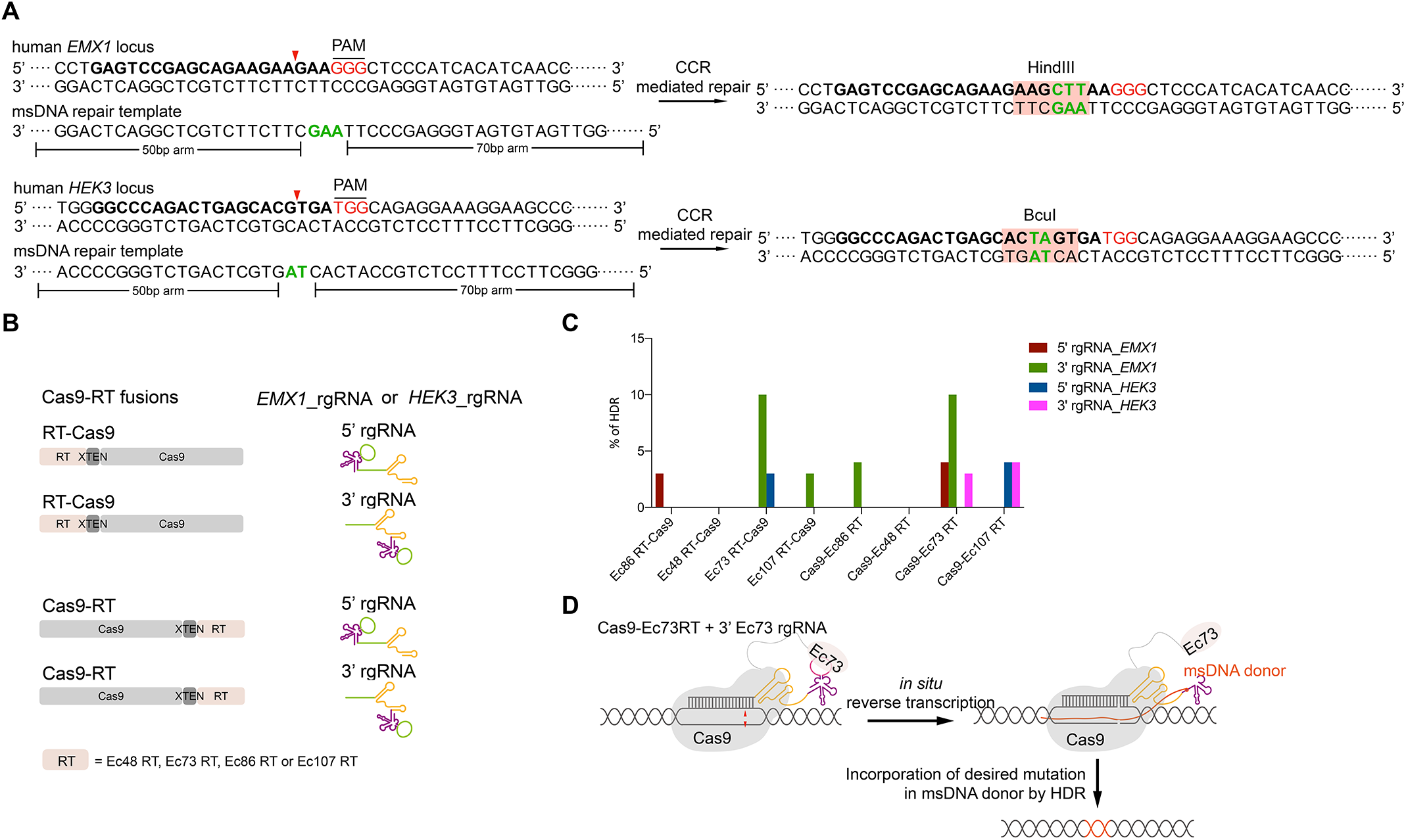
Retron editing system mediates precise editing at endogenous genomic loci. **(A)** Schematic of retron editing at human *EMX1* locus and *HEK3* locus. The reverse-transcribed msDNA repair template complementary to the non-target strand is used as a repair template. The target sequence is in bold. The insertion sequence is marked in green letter. After retron editing, HindIII and BcuI site is inserted into the *EMX1* locus and *HEK3* locus, respectively. **(B)** Plasmids construction strategies for retron editing system in human cells. Different Cas9-RT fusions targeting *EMX1* or *HEK3* combining with 5’ rgRNA or 3’ rgRNA that can produce msDNA with homology donor sequences relative to *EMX1* or *HEK3*, respectively, were co-expressed in HEK293T cells. **(C)** Blunt-end cloning assay to determine the efficiency of retron editing. Over 25 randomly selected colonies from indicated transfected cell were Sanger-sequenced. **(D)** Schematic diagram showing that Cas9-Ec73 RT fusion in-situ reverse transcribe 3’ Ec73 rgRNA to generate the msDNA containing a single-stranded donor sequence with homology arms, and msDNA is incorporated into the genome site cut by Cas9.

Of note, besides retron editing events, we detected indel containing alleles (Fig. S2B), suggesting RT-Cas9 fusions co-expressing with relevant rgRNA retained the double stranded DNA cleavage activity of Cas9.

## Discussion

The ability to write any modification of interest into the genome is a long-sought goal of biotechnology. Retrons, capable of producing intracellular ssDNA as donor with high specificity, are promising biological sources for precise genome editing. In this study, we demonstrate that different retrons are functional in human cells. Moreover, by co-expressing Cas9-RT fusions and 3’ extended retron-ncRNA gRNA, retron editing could mediate precise genome editing in human cell. Additionally, retrons coupled with CRISPR have the ability to insert a GFP gene with high efficiency in yeast (Sharon et al., 2018), making us anticipating retron editing a more versatile genome editor comparing to base editors and prime editors, which can only induce relative smaller scale genomic modifications (Anzalone et al., 2019; Gaudelli et al., 2017; Komor et al., 2016).

Many approaches may be done to improve retron editing in human cells. Firstly, identification of more suitable retron for genome editing in human cells. Second, engineered evolution of both retron-RT and retron-ncRNA can be done to enhance retron mediated intracellular production of ssDNA. Finally, evidences were shown that hRAD51 mutant fused to Cas9 (D10A) nickase (RDN) fusions could mediate precise genome editing without DSBs (Rees et al., 2019). It is plausible to anticipate that combination of retrons with RDN may make retron editing more accurate and safer due to the elimination of Cas9-induced DSBs.

To be noted, when we are preparing this manuscript, Zhao and colleagues preprinted their work of retron-mediated precise gene editing in human cells, which further convinced that retrons can be harnessed for precise genome editing in human cells by coupling with CRISPR-Cas9 (Zhao et al., 2021).

## Materials and Methods

### Plasmids construction

All DNA sequences of plasmids used in this study were provided in Supplemental Sequences. Human codon-optimized retron RTs were synthesized and inserted into EcoRI-digested pHG023 backbone or SmaI-digested pHG077 backbone by NEBuilder (New England Biolabs). The cloning of plasmids for expressing 5’ rgRNA and 3’ rgRNA were based on pHG215 backbone. Donor sequences were cloned and inserted into the replaceable region of retron ncRNA with EcoRI digestion and Gibson assembly using NEBuilder. Primers for cloning were listed in Supplemental Table1.

### Cell Culture, transfection and flow cytometry sorting

Cell Culture, transfection and flow cytometry were performed as previously described (Xu et al., 2021). Briefly, HEK293T cells cultured in DMEM supplemented with 10% FBS and penicillin/streptomycin were seeded on 24-well poly-D-lysine coated plates (Corning). Transfection was conducted with 4 μl of PEI (Polyscience) following the manufacturer’s manual and 1μg of CR plasmids and 1μg of rgRNA plasmids. Transfected cells were sorted by MoFlo XDP 72 h after transfection. BFP and mCherry double-positive cells were collected.

### msDNA purification and quantification

Four thousand sorted cells were centrifuged down at 7,000 rpm for 5 mins. Cells plate were resuspended in Trizol for isolating rgRNA-msDNA hybrid as the the manufacturer’s manual for RNA purification. For quantifying the abundance of *HEK3* msDNA, the rgRNA-msDNA extract was digested by DdeI to eliminate the contamination of plasmid DNA and genomic DNA. PCR was conducted using the DdeI digested product as template.

### Genomic DNA extraction

Four thousand sorted cells were harvested for genomic DNA extraction by addition of 10 μl of lysis buffer (Vazyme) following the manufacturer’s manual.

### Efficiency analysis of retron editing at human endogenous genomic loci

The genomic DNA extraction was performed as described above. The genomic region in the vicinity of Cas9 target site was amplified by Phanta Max Super-Fidelity DNA Polymerase (Vazyme) using nested PCR: First round: 98 °C, 3 mins; (98 °C, 30 sec; 56 °C, 30 sec; 72 °C, 1 min) X 20 cycles; 72 °C, 5 min. Second round: 98 °C, 3 mins; (98 °C, 30 sec; 52 °C, 30 sec; 72 °C, 30 sec) X 35 cycles; 72 °C, 5 min. All primers used for genomic DNA amplification are listed in Supplemental Table1.

Purified PCR products were inserted into TA/Blunt-Zero Cloning vector (Vazyme). Plasmids were isolated from randomly selected colonies and subjected for Sanger sequencing. The retron editing frequency was calculated as the ratio of number of colony containing precise editing vs. total sequenced colony number.

Restriction-fragment length polymorphism (RFLP) was performed to determine the CR-mediated precise editing. Purified PCR products were digested with HindIII and BcuI (Thermo) for *EMX1* and *HEK3* amplicon respectively, and incubated for 90 mins at 37 °C. The digested products were analyzed by agarose gel electrophoresis and imaged with gel imaging system (Tanon).

## Acknowledgments

This work was supported by HUIGENE Therapeutics Inc., the Basic Frontier Scientific Research Program of Chinese Academy of Sciences From 0 to 1 original innovation project (grant no. ZDBS-LY-SM001), the R&D Program of China (grant nos. 2017YFC1001300 and 2018YFC2000100) and the CAS Strategic Priority Research Program (grant no. XDB32060000),

## Authors’ Contributions

X.K. and H.Y. conceived and designed the project. X.K., Z.W. and X.W. performed experiments and analyzed data. Y.Z. analyzed the data. X.K., L.S. and H.Y. wrote the manuscript.

---

**Supplemental Figure 1.**
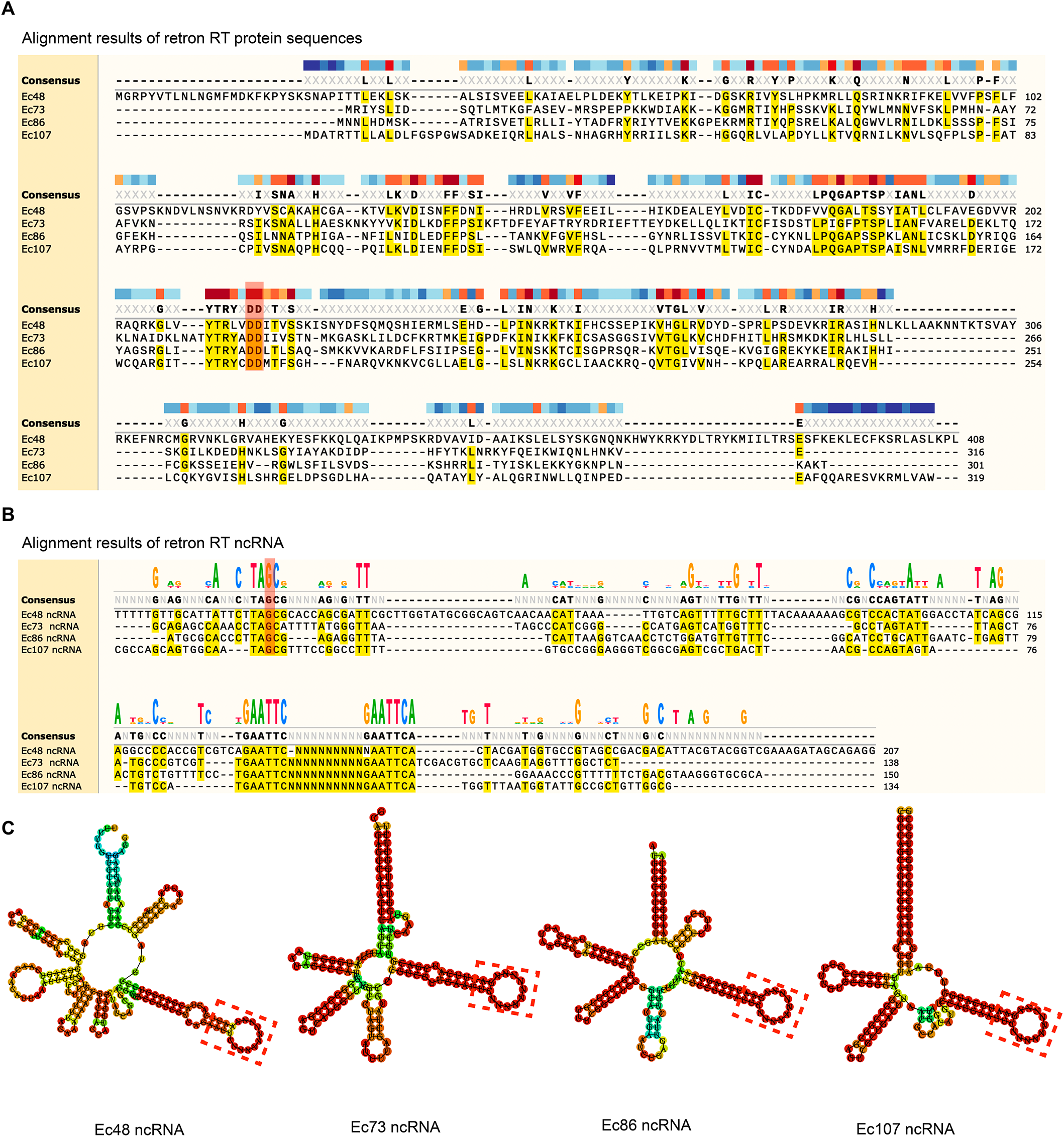
Comparison of four retrons. **(A)** Protein sequences alignment of Ec48-RT, Ec73-RT, Ec86-RT and Ec107-RT. Conserved double aspartic acid (DD) were highlighted in red. **(B)** Alignment of Ec48-ncRNA, Ec73-ncRNA, Ec86-ncRNA and Ec107-ncRNA. Conserved priming G was highlighted in red. **(C)** Predicted secondary structure of Ec48-ncRNA, Ec73-ncRNA, Ec86-ncRNA and Ec107-ncRNA using RNAfold web server. The replaceable region of retron ncRNA was marked with red dotted line.

**Supplemental Figure 2.**
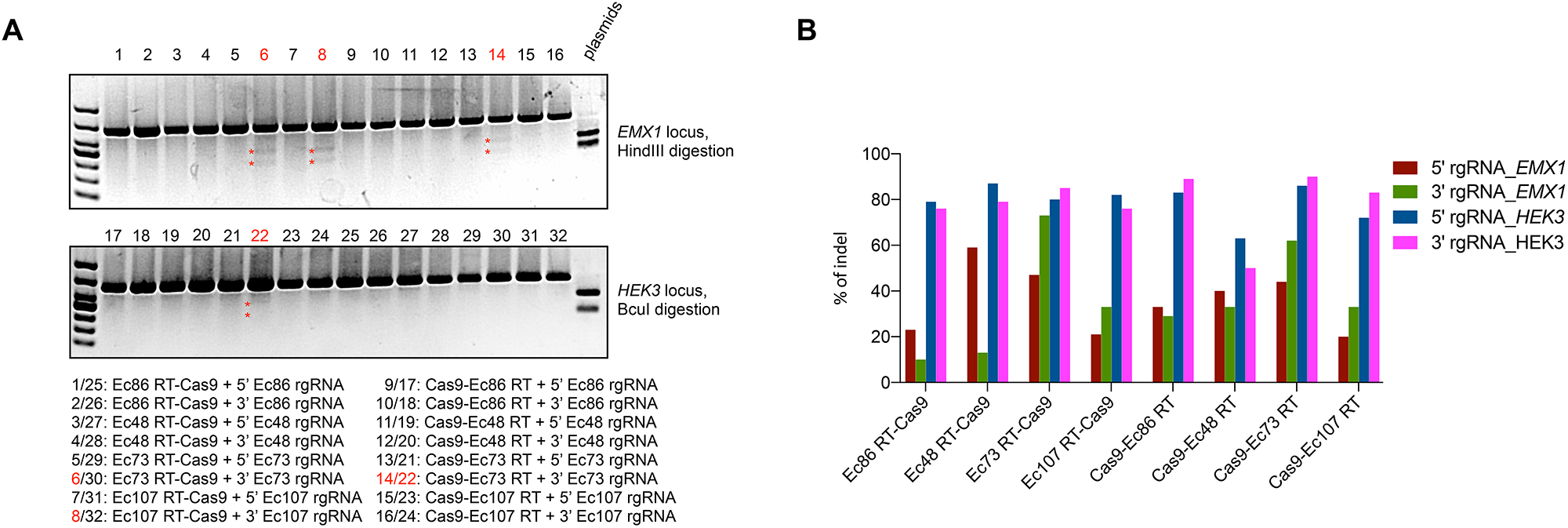
Retron editing system-stimulated precise editing and indels. **(A)** Determination of retron editing system-mediated precise genome editing by restriction-fragment length polymorphism (RFLP) analysis. Digested products were marked by asterisks. **(B)** Analysis of retron editing system-induced indels.

**Supplemental Sequences. Cas9-RTs fusion plasmids and rgRNA expression plasmids.**

**Supplemental Table1.**
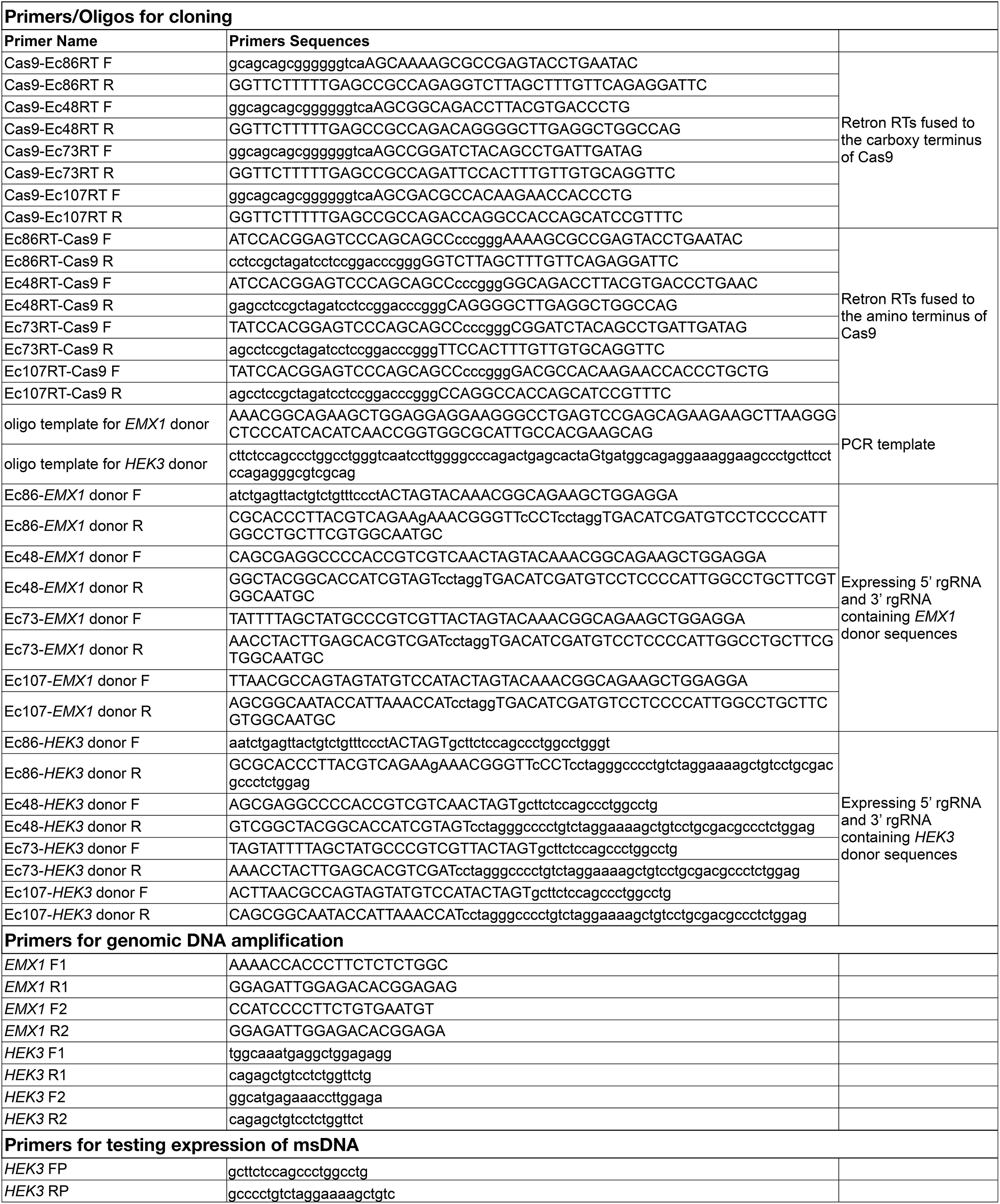
Primers and oligos used in this study.

## Supplemental Sequences

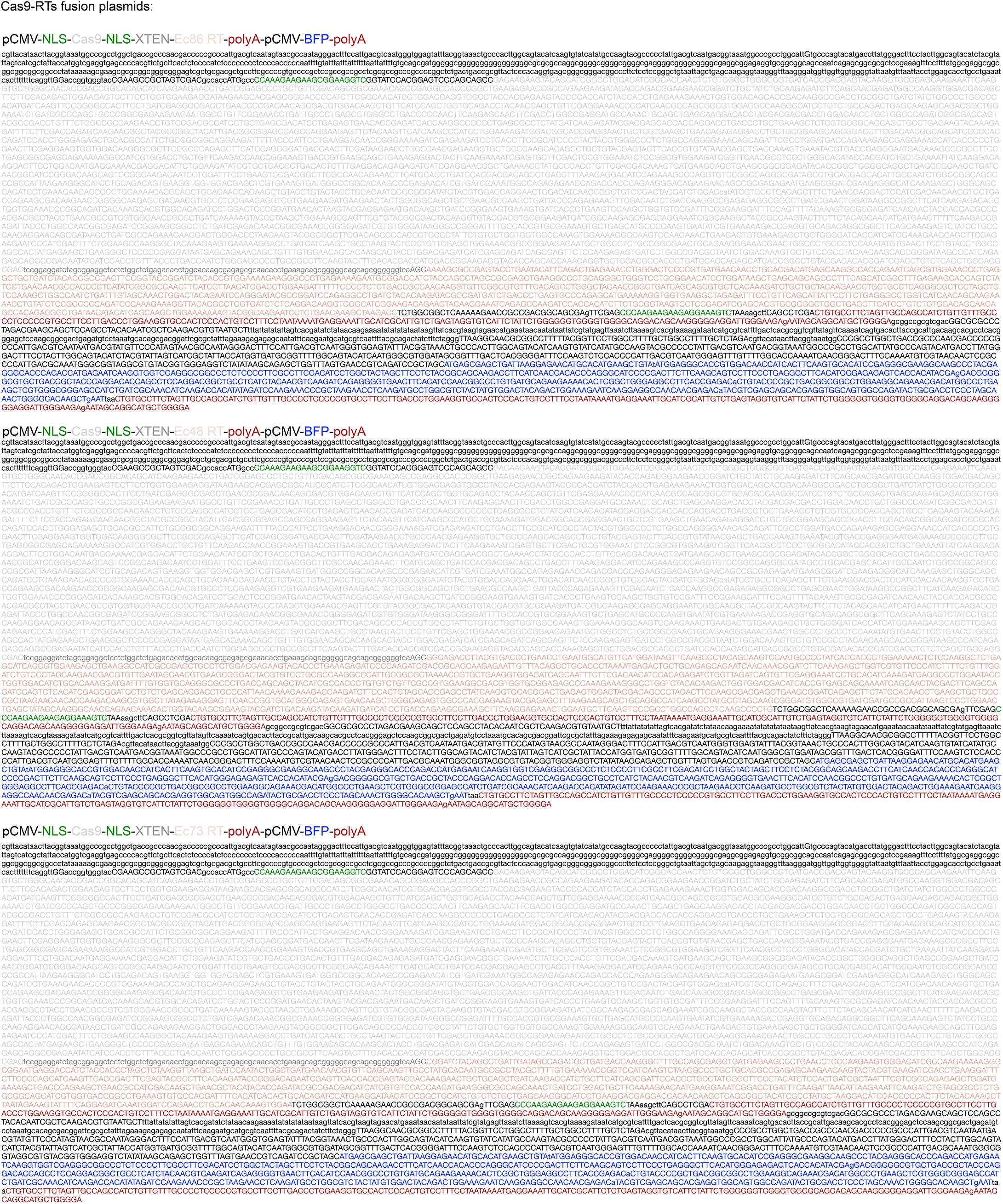
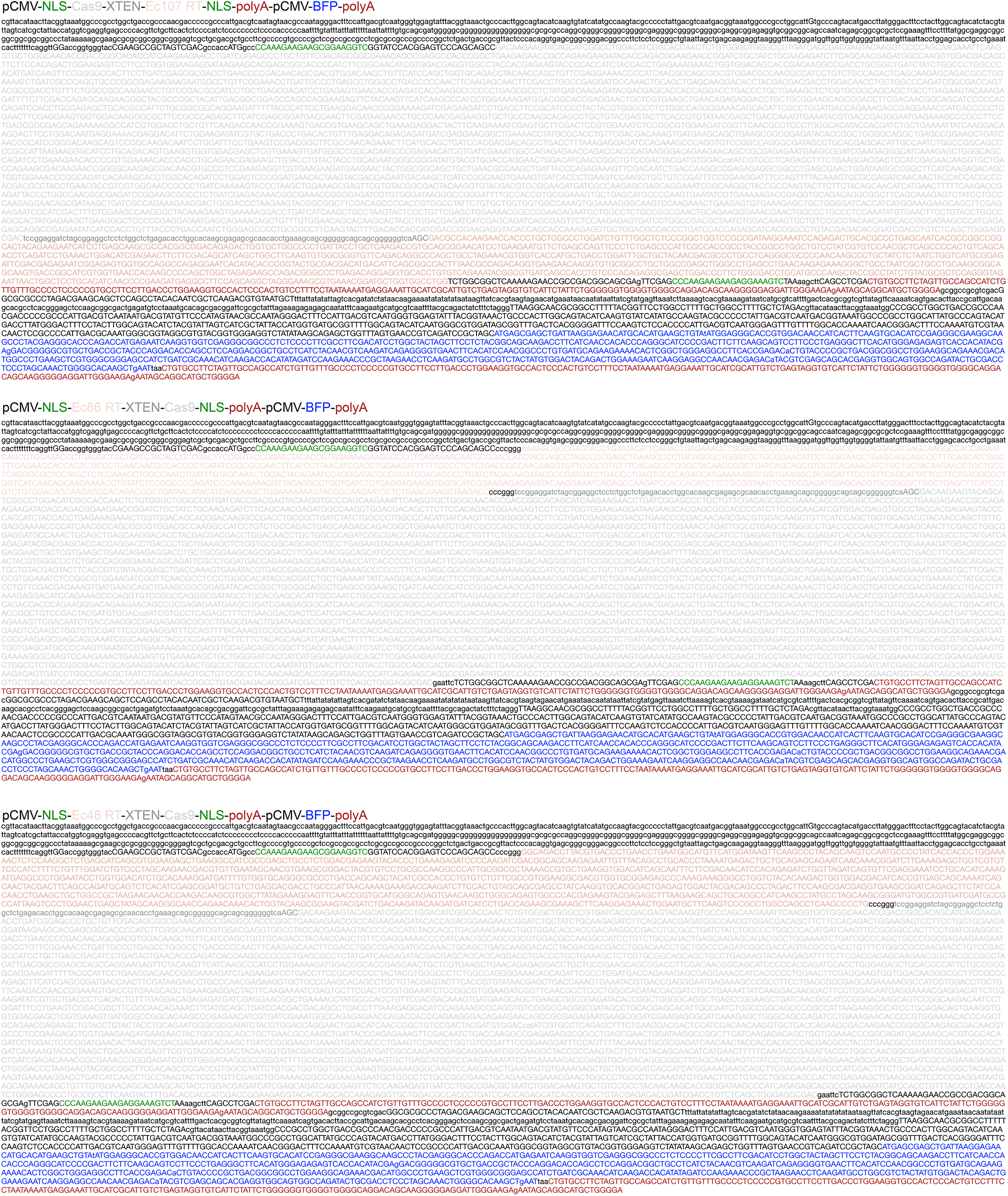
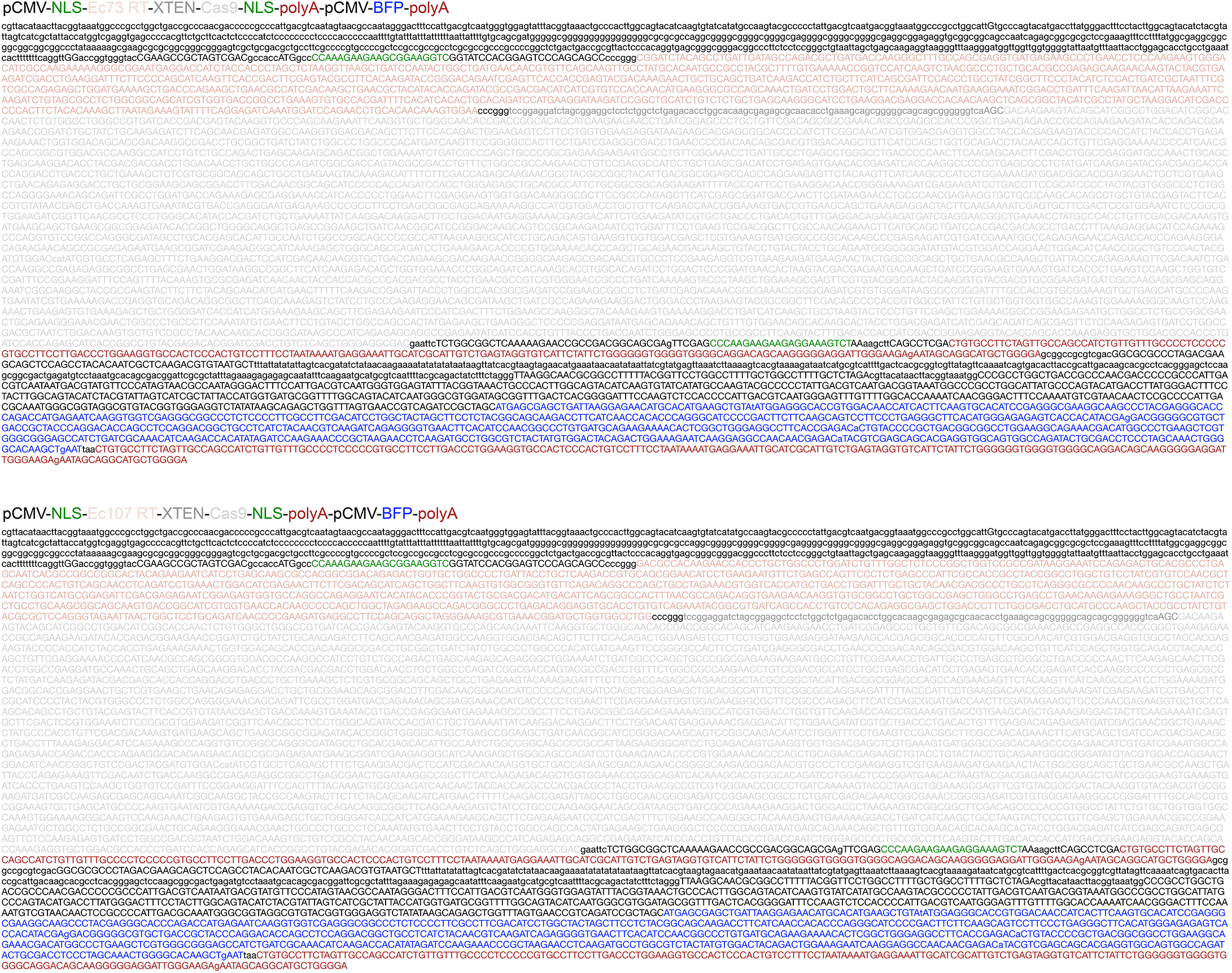

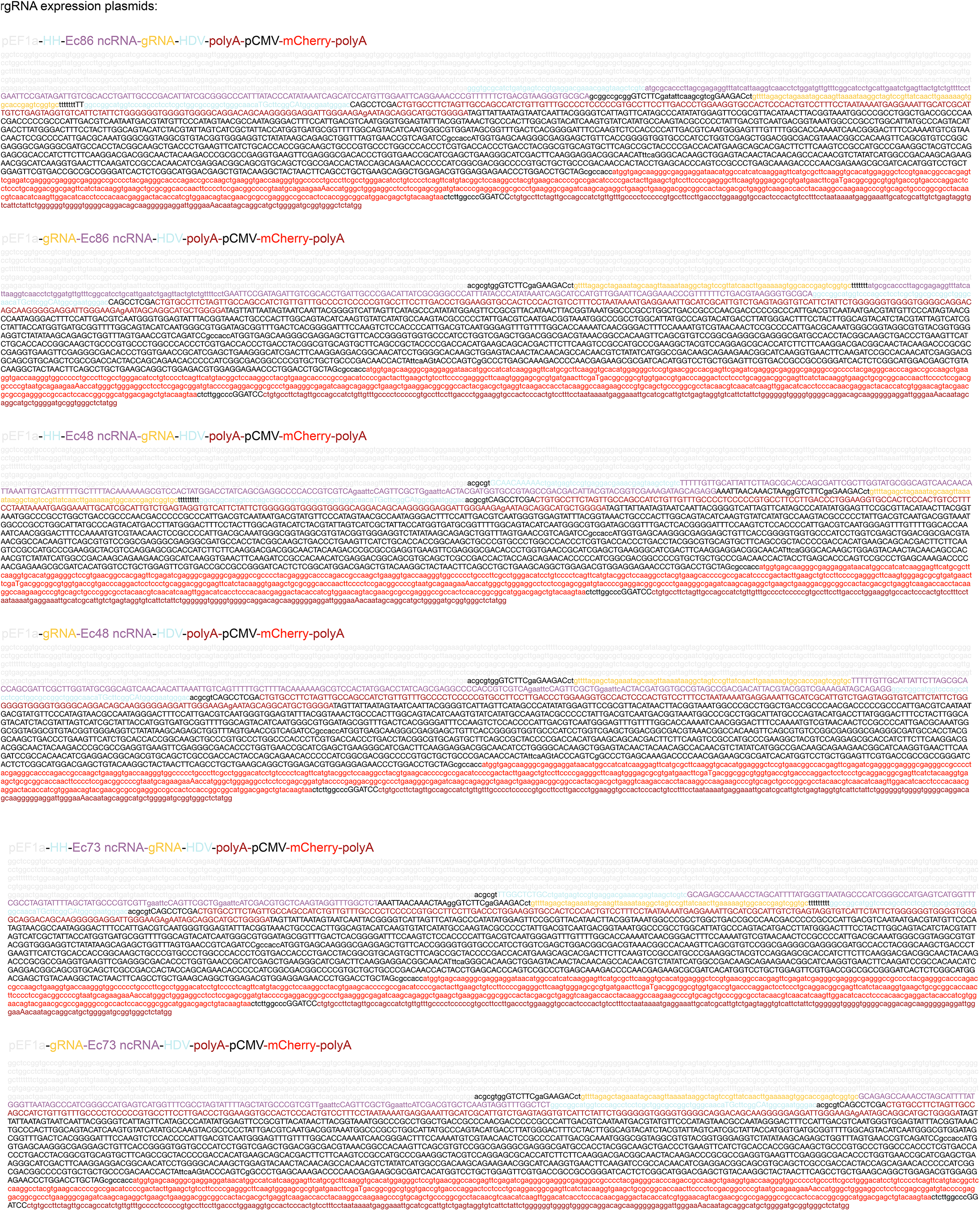
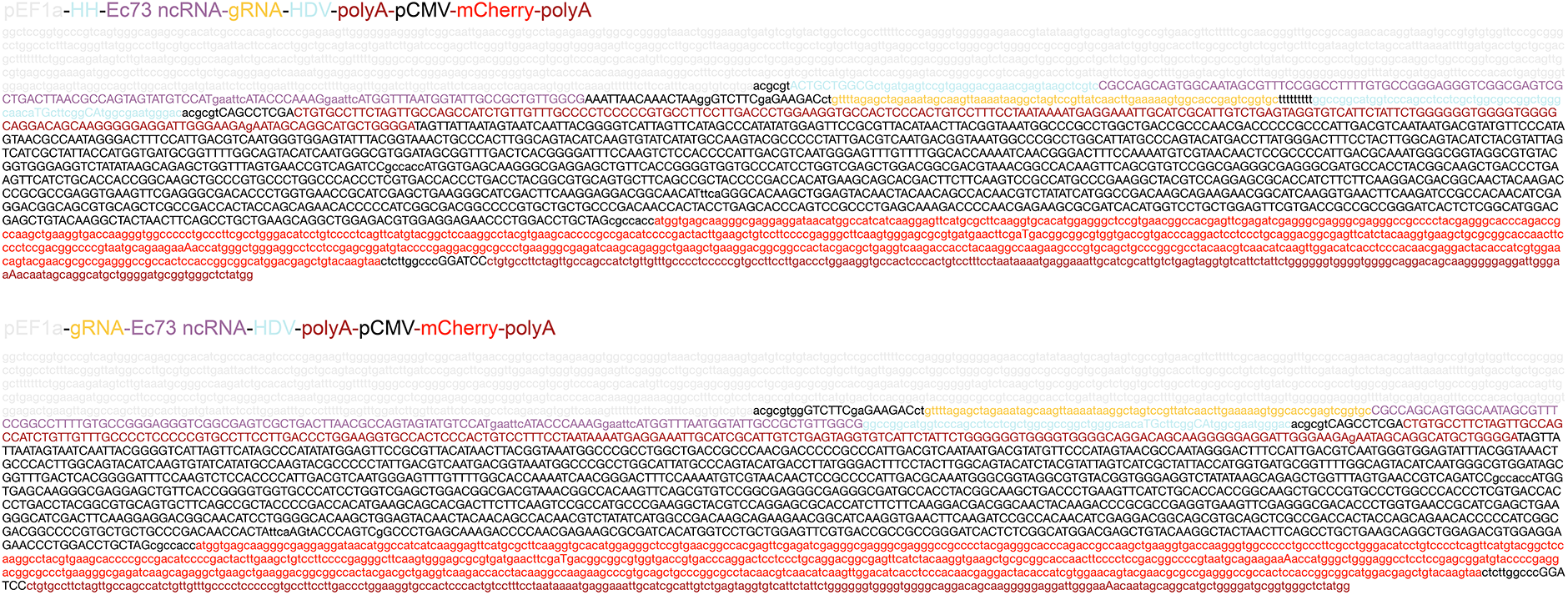

## References

Aird, E.J., Lovendahl, K.N., St Martin, A., Harris, R.S., and Gordon, W.R. (2018). Increasing Cas9-mediated homology-directed repair efficiency through covalent tethering of DNA repair template. Commun Biol 1, 54.

Anzalone, A.V., Koblan, L.W., and Liu, D.R. (2020). Genome editing with CRISPR-Cas nucleases, base editors, transposases and prime editors. Nat Biotechnol 38, 824–844.

Anzalone, A.V., Randolph, P.B., Davis, J.R., Sousa, A.A., Koblan, L.W., Levy, J.M., Chen, P.J., Wilson, C., Newby, G.A., Raguram, A., et al. (2019). Search-and-replace genome editing without double-strand breaks or donor DNA. Nature 576, 149-+.

Chu, V.T., Weber, T., Wefers, B., Wurst, W., Sander, S., Rajewsky, K., and Kuhn, R. (2015). Increasing the efficiency of homology-directed repair for CRISPR-Cas9-induced precise gene editing in mammalian cells. Nat Biotechnol 33, 543–548.

Cong, L., Ran, F.A., Cox, D., Lin, S., Barretto, R., Habib, N., Hsu, P.D., Wu, X., Jiang, W., Marraffini, L.A., et al. (2013). Multiplex genome engineering using CRISPR/Cas systems. Science 339, 819–823.

Cox, D.B., Platt, R.J., and Zhang, F. (2015). Therapeutic genome editing: prospects and challenges. Nat Med 21, 121–131.

Dhundale, A., Lampson, B., Furuichi, T., Inouye, M., and Inouye, S. (1987). Structure of msDNA from Myxococcus xanthus: evidence for a long, self-annealing RNA precursor for the covalently linked, branched RNA. Cell 51, 1105–1112.

Doudna, J.A. (2020). The promise and challenge of therapeutic genome editing. Nature 578, 229–236.

Farzadfard, F., and Lu, T.K. (2014). Synthetic biology. Genomically encoded analog memory with precise in vivo DNA writing in living cell populations. Science 346, 1256272.

Gao, C. (2021). Genome engineering for crop improvement and future agriculture. Cell 184, 1621–1635.

Gao, L., Altae-Tran, H., Bohning, F., Makarova, K.S., Segel, M., Schmid-Burgk, J.L., Koob, J., Wolf, Y.I., Koonin, E.V., and Zhang, F. (2020). Diverse enzymatic activities mediate antiviral immunity in prokaryotes. Science 369, 1077–1084.

Gaudelli, N.M., Komor, A.C., Rees, H.A., Packer, M.S., Badran, A.H., Bryson, D.I., and Liu, D.R. (2017). Programmable base editing of A*T to G*C in genomic DNA without DNA cleavage. Nature 551, 464–471.

Hsu, M.Y., Inouye, S., and Inouye, M. (1989). Structural requirements of the RNA precursor for the biosynthesis of the branched RNA-linked multicopy single-stranded DNA of Myxococcus xanthus. J Biol Chem 264, 6214–6219.

Inouye, M., Mao, J.R., Shimamoto, T., and Inouye, S. (1997). In vivo production of oligodeoxyribonucleotides of specific sequences: application to antisense DNA. Ciba Found Symp 209, 224–233; discussion 233-224.

Komor, A.C., Kim, Y.B., Packer, M.S., Zuris, J.A., and Liu, D.R. (2016). Programmable editing of a target base in genomic DNA without double-stranded DNA cleavage. Nature 533, 420–424.

Lampson, B.C., Inouye, M., and Inouye, S. (1989). Reverse transcriptase with concomitant ribonuclease H activity in the cell-free synthesis of branched RNA-linked msDNA of Myxococcus xanthus. Cell 56, 701–707.

Lim, H., Jun, S., Park, M., Lim, J., Jeong, J., Lee, J.H., and Bang, D. (2020). Multiplex Generation, Tracking, and Functional Screening of Substitution Mutants Using a CRISPR/Retron System. ACS Synth Biol 9, 1003–1009.

Lin, S., Staahl, B.T., Alla, R.K., and Doudna, J.A. (2014). Enhanced homology-directed human genome engineering by controlled timing of CRISPR/Cas9 delivery. Elife 3, e04766.

Lopez, S.C., Crawford, K.D., Bhattarai-Kline, S. and Shipman, S.L. (2021). Improved architectures for flexible DNA production using retrons across kingdoms of life. BioRxiv preprint doi: https://doi.org/10.1101/2021.03.26.437017.

Ma, M., Zhuang, F., Hu, X., Wang, B., Wen, X.Z., Ji, J.F., and Xi, J.J. (2017). Efficient generation of mice carrying homozygous double-floxp alleles using the Cas9-Avidin/Biotin-donor DNA system. Cell Res 27, 578–581.

Ma, S., Wang, X., Hu, Y., Lv, J., Liu, C., Liao, K., Guo, X., Wang, D., Lin, Y., and Rong, Z. (2020). Enhancing site-specific DNA integration by a Cas9 nuclease fused with a DNA donor-binding domain. Nucleic Acids Res 48, 10590–10601.

Mao, J.R., Shimada, M., Inouye, S., and Inouye, M. (1995). Gene regulation by antisense DNA produced in vivo. J Biol Chem 270, 19684–19687.

Mao, Y.F., Botella, J.R., Liu, Y.G., and Zhu, J.K. (2019). Gene editing in plants: progress and challenges. Natl Sci Rev 6, 421–437.

Maruyama, T., Dougan, S.K., Truttmann, M.C., Bilate, A.M., Ingram, J.R., and Ploegh, H.L. (2015). Increasing the efficiency of precise genome editing with CRISPR-Cas9 by inhibition of nonhomologous end joining. Nat Biotechnol 33, 538–542.

Millman, A., Bernheim, A., Stokar-Avihail, A., Fedorenko, T., Voichek, M., Leavitt, A., Oppenheimer-Shaanan, Y., and Sorek, R. (2020). Bacterial Retrons Function In Anti-Phage Defense. Cell 183, 1551–1561 e1512.

Mirochnitchenko, O., Inouye, S., and Inouye, M. (1994). Production of single-stranded DNA in mammalian cells by means of a bacterial retron. J Biol Chem 269, 2380–2383.

Paquet, D., Kwart, D., Chen, A., Sproul, A., Jacob, S., Teo, S., Olsen, K.M., Gregg, A., Noggle, S., and Tessier-Lavigne, M. (2016). Efficient introduction of specific homozygous and heterozygous mutations using CRISPR/Cas9. Nature 533, 125–129.

Rees, H.A., Yeh, W.H., and Liu, D.R. (2019). Development of hRad51-Cas9 nickase fusions that mediate HDR without double-stranded breaks. Nat Commun 10, 2212.

Richardson, C.D., Ray, G.J., DeWitt, M.A., Curie, G.L., and Corn, J.E. (2016). Enhancing homology-directed genome editing by catalytically active and inactive CRISPR-Cas9 using asymmetric donor DNA. Nature Biotechnology 34, 339-+.

Savic, N., Ringnalda, F.C., Lindsay, H., Berk, C., Bargsten, K., Li, Y., Neri, D., Robinson, M.D., Ciaudo, C., Hall, J., et al. (2018). Covalent linkage of the DNA repair template to the CRISPR-Cas9 nuclease enhances homology-directed repair. Elife 7.

Schubert, M.G., Goodman, D.B., Wannier, T.M., Kaur, D., Farzadfard, F., Lu, T.K., Shipman, S.L., and Church, G.M. (2021). High-throughput functional variant screens via in vivo production of single-stranded DNA. Proc Natl Acad Sci U S A 118.

Sharon, E., Chen, S.A., Khosla, N.M., Smith, J.D., Pritchard, J.K., and Fraser, H.B. (2018). Functional Genetic Variants Revealed by Massively Parallel Precise Genome Editing. Cell 175, 544–557 e516.

Yang, H., Wang, H., Shivalila, C.S., Cheng, A.W., Shi, L., and Jaenisch, R. (2013). One-step generation of mice carrying reporter and conditional alleles by CRISPR/Cas-mediated genome engineering. Cell 154, 1370–1379.

Yee, T., Furuichi, T., Inouye, S., and Inouye, M. (1984). Multicopy single-stranded DNA isolated from a gram-negative bacterium, Myxococcus xanthus. Cell 38, 203–209.

Zhao, B., Chen S-A. A., Lee J, Fraser H.B. (2021). Bacterial retrons enable precise gene editing in human cells. BioRxiv preprint doi: https://doi.org/10.1101/2021.03.29.437260

